# Non-disclosure of tuberculosis diagnosis by patients to their household members in south western Uganda

**DOI:** 10.1101/622597

**Authors:** Miria Nyangoma, Francis Bajunirwe, Daniel Atwine

**Affiliations:** Faculty of Computing and Informatics, Mbarara University of Science and Technology, P.O. BOX 1410, Mbarara, Uganda; Department of Community Health, Mbarara University of Science and Technology, P.O.BOX 1410, Mbarara, Uganda; Epicentre, Mbarara, P.O BOX 1956, Mbarara, Uganda

**Keywords:** tuberculosis, disclosure, stigma, infection control, gender, religion

## Abstract

**Background:** Tuberculosis (TB) non-disclosure by adult patients to all household members is a setback to TB control efforts. It reduces the likelihood that household contacts will seek early TB screening, initiation on preventive or curative treatment, but also hinders the implementation of infection controls and home-based directly observed treatment. Therefore, this study aimed at determining the level of TB non-disclosure, its predictors and effects of disclosure to household members on adult patients at a large regional referral hospital in south-western Uganda.

**Methods:** Cross-sectional study. Questionnaires administered to collect patients’ sociodemographic and their TB disclosure data. Non-disclosure was considered if a patient did not reveal their TB diagnosis to all household members within 2 weeks post-treatment initiation. Univariate and multivariate logistic regression models were fitted for predictors of non-disclosure.

**Results:** Enrolled 62 patients, 74% males, mean age of 32 years, and median of five people per household. Non-disclosure rate was 30.6%. Post-disclosure experiences were positive in 98.3% of patients, while negative experiences suggestive of severe stigma occurred in 24.6% of patients. Being female (OR 6.5, 95% CI: 1.42-29.28) and belonging to Muslim faith (OR 12.4, 95% CI: 1.42-109.05) predicted TB non-disclosure to household members.

**Conclusions:** There is a high rate of TB non-disclosure to household members by adult patients in rural Uganda, with the highest vulnerability seen among female and Muslim patients. Interventions enhancing TB disclosure at household level while minimizing negative effects of stigma should be developed and prioritized.

## BACKGROUND

Tuberculosis (TB) remain as one of the top 10 causes of death worldwide and the leading cause from a single infectious agent, ranking above HIV/AIDS [1]. The world health organization (WHO) estimated that 10 million people fell ill with TB in 2017 and that treatment success rate among new cases remained at only 82% globally [1]. Uganda is among the top 30 TB-HIV high burden countries [1], with an overall TB prevalence of 253 per 100,000 people and incidence of 234 per 100,000 people [2].

In addition to the HIV epidemic, uncontrolled transmission is fueling the global tuberculosis epidemic [3]. All indoor environments especially homes and congregate settings are potential sites of transmission since air dilution is limited and occupants are concentrated [4–5]. Ensuring patient adherence to effective treatment guarantees a faster reduction in risk of TB transmission from adult patients to their household members [6], but also it limits the emergence of drug resistance. In this regard, WHO recommends both social and psychological support as supplements to community- or home-based directly observed treatment [7]. This would ensure treatment effectiveness; prevent death from active TB or its late effects; prevent TB relapse; but also reduce transmission to others [8]; and prevent the development and transmission of drug resistance [9–10]. Noteworthy, household members are well positioned to provide such needed social and psychological support so as to ensure good treatment outcomes and infection control [11], but only if adequately engaged in care of their TB patient, a step that is entirely dependent on patients’ disclosure of their TB diagnosis.

Patients’ disclosure of their TB diagnosis to household members could influence their treatment adherence [12], but also allow early initiation of appropriate infection control measures, and access to diagnostic, curative and preventive TB services by their household contacts. TB disclosure, however can be hindered in the presence of TB-related stigma, which has been associated with human immunodeficiency virus (HIV)-infection, alleged immoral behavior, perceived incurability of TB, and myths about TB etiology [13]. Negative consequences of disclosure like patient isolation, neglect, cut-off of support, and divorce have been previously reported [14]. It is therefore important to understand these negative consequences as potential barriers to TB disclosure and to develop remedies to overcome them.

However, there is a paucity of information documented on the TB disclosure at household level and its impact on patients within TB-HIV high-burden countries, including Uganda. Such information is useful in designing contextualized interventions to promote disclosure, as part of the efforts to achieve the 2035 targets of the End-TB strategy[15]. Overall, knowledge on TB disclosure dynamics at household level, could guide TB control interventions such as home-based directly observed treatment, short-course (DOTS), also known as TB-DOTS, TB contact screening with initiation of preventive therapy, and TB infection control at household level, whose success is dependent on household members’ involvement in patient care. Therefore, we conducted a study at a large regional referral hospital in south-western Uganda, aimed at measuring the level of TB non-disclosure to household members by adult patients, its predictors and patients’ experiences post- disclosure.

## METHODS

### Design and setting

We conducted a cross sectional study among adult patients started on antituberculosis treatment for pulmonary TB at the TB clinic at Mbarara Regional Referral Hospital, a large tertiary health facility in south-western Uganda with a catchment area of nearly 2 million people. Data were collected at the clinic between May and July 2017. In routine practice, in agreement with the National TB Control Program guidelines, prior to initiation of TB treatment, each patient is required to identify someone close to them, to serve as a treatment supporter where possible. Trained nurses then counsel patients on TB disease, treatment adherence and infection control before administering treatment. Patients return for drug refills every 2 weeks during the first 2 months of intensive phase, and thereafter, monthly during the 4-month continuation phase.

### Study population

Patients were eligible for study participation if they were 18 years or older, either new or retreatment case with bacteriologically confirmed or unconfirmed pulmonary TB, completed at least 2 weeks of TB treatment, having household members, and with written consent. In this study, the household was taken to consist of more than one person living in the same dwelling, share meals, living accommodation and may consist of a single family or some other grouping of people.

Patients were excluded if they were too ill, or had psychiatric disorders deterring them from responding to interview questions. The informed consent process was administered by trained study nurses that were not participating in patient care within the TB clinic.

### Study procedures

The study utilized only quantitative data collection methods. Structured questionnaires with both closed and open questions were administered by study nurses in the local language or English, as per the patient’s preference. The collected information included; socio-demographic, household, patients’ behavioral factors especially alcohol intake and smoking, medical history, and patients’ TB disclosure experiences. We focused at assessing early disclosure, which we defined in this study as patients’ TB disclosure to household members within the first 2 weeks following treatment initiation. The choice of the first 2 weeks was premised on it being the estimated time for a patient on appropriate TB treatment to become non-infectious [16], and therefore, could serve as the best time to optimize adherence support but also infection control measures so as to minimize TB transmission at household level.

In addition, post-disclosure experiences were elicited by asking patients both closed and open questions on their experiences following disclosure to household members, and the perceived impact of such experiences on them. With respect to positive experiences, questions were aligned towards establishing any reception of psychosocial support and support related to enhancing patients’ treatment adherence and compliance to scheduled follow-up visits to clinic. Patients were asked to respond “yes” or “No” on whether they experienced any of the following after disclosing their TB diagnosis to household members, that is, whether they received; 1) encouragement from family members, 2) support in taking their medications, 3) support in feeding, 4) financial support to attend clinic days, 5) Other support, with an additional open question for them to specify the other support received. In the same way, closed and open questions were asked to elicit the negative experiences patients faced after TB disclosure to household members. These questions were focused on demonstrating any effects on the patient related to potential stigma related to TB among household members. These included experiences of; 1) criticism or blame, 2) isolation by household members, 3) withdrawal of support by household, 4) marital separation, and 5) other negative experiences, with an additional open question for them to specify these other negative experiences faced.Patients were then asked to respond “yes” or “No” on whether they perceived that the above positive or negative post-disclosure experiences had impacted on their treatment intake and compliance to scheduled TB clinic visits, but also on some of their psychosocial aspects, that is; self-esteem, hope of recovery, work, household harmony, and marital relationships in the case of those who were married or co-habiting. Also, patients were asked on whether they would agree to the training of their treatment supporters by healthcare workers so as to support in patients’ TB disclosure to household members.

The dependent variable for this study was “disclosure”, which was derived from a question which asked the patients whether they had “disclosed their current TB status to their household members”, with three responses — “yes, all = 1”, “yes, some = 2” and “no = 3”. A patient was considered to have complete disclosure if he or she responded with “yes, all”, meaning that they had revealed their TB diagnosis to all their household members. A patient was considered to have partial disclosure if he or she responded with “yes, some”, meaning that he or she had revealed their TB diagnosis to only a selected number of their household members and left out others.

For the purpose of the predictor analysis, we generated a binary dependent variable coded 0= complete disclosure, and 1= non-disclosure. The “no disclosure’ category here, was derived by grouping both partial disclosure and complete non-disclosure. The decision was made because both total non-disclosure and partial disclosure place household members at risk of TB infection, given that TB is an infectious disease. The independent variables included all patients’ characteristics, that is, socio-demographics, household characteristics, patients’ behavioural factors especially alcohol intake and smoking and medical characteristics.

### Sample size

A sample size of 62 patients was calculated using a formula for single population proportion with correction for finite population [17], that is: n = N*X / (X + N − 1), where, X = Z_α/2_^2^ *p*(1-p) / MOE^2^, and Z_α/2_ is the critical value of the Normal distribution at α/2 for α of 0.05, MOE is the margin of error considered at 5%, p is the proportion of non-disclosure which was taken as 50%, an arbitrary value given the lack of comparable data, and N is the population size, estimated at 70 new patients with TB undergoing treatment at the clinic for two weeks to two month period. To cater for attrition, an additional 5% was added to the sample size.

### Data Analysis

Data from completed questionnaires were entered into a database designed using Epi Info™ software (V7.2, 1600 Clifton Road Atlanta, GA 30329-4027 USA) and analysis was performed with Stata software (v.13, College Station, Texas, USA). Descriptive statistics were generated for participants’ characteristics. The proportion of patients reporting not to have disclosed their TB status (that is, partially or completely) to household members was calculated out of all study participants and presented as a percentage. Percentages of TB patients with positive or negative post-disclosure experiences, and those reporting different aspects of their lives impacted by these post-disclosure experiences at household level were calculated and presented as bar-graphs. The dependent variable was a binary variable of TB status non-disclosure to household members coded 0=No and 1=Yes. Univariate and multivariate analysis using logistic regression was performed to establish the factors associated with TB status non-disclosure to household members. A significance level of 5% was maintained. Both unadjusted and adjusted odds ratios with corresponding 95% confidence intervals were reported. Percentages were calculated to describe the patients’ perception on training treatment supporters so as to enhance their role in supporting patients’ disclosure at household level.

### Role of funding source

The funder of the study had no role in study design, data collection, analysis, interpretation, or writing of the report. The first and last authors had full access to all data and had final responsibility for the decision to submit the manuscript for publication.

The study was approved by the Mbarara University Faculty of Medicine Research Committee, and Mbarara University of Science and Technology Research Ethics Committee. Study approval no: 16/02-17. Written informed consent was received from all study participants prior to enrollment in the study. We also used separate research assistants (nurses) who were not involved in the routine care of these TB patients who interviewed patients after they had received their routine care, so as to prevent coercion and therapeutic misconception. No incentives were given for patients’ participation in the study. All patients’ information was kept confidential.

## RESULTS

We enrolled 62 patients, predominantly males (74%), mean age of 32 years, with half having at least secondary education. The median household size was 5 (range: 1-22) people, with 80% of the households having at least four people. About 95% of the patients had the current episode as their first TB episode, 82% had a treatment supporter, 49 % HIV infected, and two-thirds had been on TB treatment for at least 8 weeks at the time of interview. The details of the sociodemographic characteristics are shown in Table 1.

**Table 1.** Participants’ characteristics

| Characteristics | N | n (%) |
| --- | --- | --- |
| Age in years, mean (SD) | 62 | 32.3 (8.6) |
| Age categories, n (%) | 62 |  |
| <25 |  | 12 (19.4) |
| 25-34 |  | 27 (43.6) |
| 35-59 |  | 23 (37.1) |
| Gender, Males, n (%) | 62 | 46 (74.2) |
| Residence, Urban, n (%) | 62 | 36 (58.6) |
| Education Level, n (%) | 62 |  |
| None |  | 8 (13.0) |
| Primary |  | 23 (37.1) |
| Secondary |  | 24 (38.7) |
| Tertiary |  | 7 (11.3) |
| Marital status, single, n (%) | 61 | 34 (55.7) |
| Occupation, n (%) | 62 |  |
| Un employed |  | 6 (9.7) |
| Business |  | 23 (37.1) |
| Peasant farmer |  | 13 (21.1) |
| Professional |  | 15 (24.2) |
| Family, n (%) | 62 |  |
| Extended |  | 15(24.2) |
| Nuclear |  | 42 (67.7) |
| Single parent |  | 4 (6.5) |
| Religion | 62 |  |
| Christian |  | 55 (88.7) |
| Moslem |  | 7 (11.3) |
| Household members, $\geq 4$, n (%) | 61 | 49 (80.3) |
| First TB episode, n (%) | 62 | 59 (95.0) |
| Treatment duration at the time of data collection, | 62 |  |
| Median (IQR) |  | 12.4 (6.4-21.3) |
| >8weeks, n (%)* |  | 40 (65.6) |
| Treatment supporter, n (%) | 62 | 51 (82.3) |
| Monthly income in Ug.sh, median (IQR) | 47 | 100,000 (50,000-200,000) |
| HIV positive, n (%) | 62 | 30 (48.4) |
| Has dependants, n (%) | 62 | 39 (62.9) |
| Patient is household head, n (%) | 62 | 31 (50.0) |
| Alcohol history prior to TB, n (%) | 62 | 39 (62.9) |
| Currently Smoking, n (%) | 62 | 16 (25.8) |

\*1 missing response. IQR, interquartile range; SD, standard deviation; TB, tuberculosis; Ug.sh, Ugandan shillings

### TB non-disclosure to household members

Overall, of the 62 patients, 19 (30.6%) had either not disclosed to some or all the members of their household. Significantly higher TB non-disclosure rates were noted among patients who were; female (62.5%, p=0.001), single (44.1%, p=0.005), and moslems (71.4%, p=0.013), as compared to their counterparts. No significant differences in non-disclosure rates were observed across patients’ age, occupation, education and residence type, p>0.05. The detail of these results are shown in Table 2.

**Table 2.** TB Non-disclosure rates across patients’ sociodemographic characteristics

| Non-disclosure rate | n/N | % [95% CI] | p value |
| --- | --- | --- | --- |
| Overall rate | 19/62 | 30.7 [20.2-43.5] |  |
| Gender-specific rates |  |  | 0.001 |
| Male | 9/46 | 19.6 [10.2-34.1] |  |
| Female | 10/16 | 62.5 [34.8-83.9] |  |
| Age-specific rates |  |  | 0.492 |
| <25 | 4/12 | 33.3 [10.9-67.1] |  |
| 25-34 | 10/27 | 37.0 [20.3-57.5] |  |
| 35-59 | 5/23 | 21.7 [8.7-44.8] |  |
| Marital status |  |  | 0.005 |
| Single | 15/34 | 44.1 [27.9-61.7] |  |
| Married | 3/27 | 11.1 [3.3-31.1] |  |
| Religion |  |  | 0.013 |
| Christian | 14/55 | 25.5 [15.4-39.0] |  |
| Moslem | 5/7 | 71.4 [21.5-95.8] |  |

CI: confidence interval

Overall, the median duration on TB treatment for patients that had not disclosed to household members by time of interview was 13.3 (IQR 2.0-13.2) weeks.

Of the five patients that had not disclosed to any one, four had been on treatment for more than 4 weeks at the time of interview.

### Motivation for TB disclosure

Of the 57 patients with either complete or partial disclosure, 50 responded to the open-ended question on what had motivated their TB disclosure to household members. The responses were, 1) The desire to receive support/care from household members (40%); 2) need to avoid transmitting the disease (30%); 3) Need for support to collect and remind them to take their medicines (16%); and then, 4) the need to inform their family what they were suffering from (14%).

### Factors associated with TB non-disclosure

Factors significantly associated with adult TB non-disclosure to household members are shown in Table 3 and were; female gender and being a Moslem, after controlling for age of patients at multivariate analysis. The odds of non-disclosure of TB status to household members were 6.5 times higher in females as compared to males (OR 6.5, [95%CI: 1.42-21.2], p=0.016). Moslem patients had 12.4 times higher odds of non-disclosure of their TB status to household members as compared to Catholic patients (OR 12.4, [95% CI: 1.42-109.05], p=0.023). Patients who were not married (single) had 4.6 times higher odds of non-disclosure of TB status to household members as compared to patients who were married, with a tendency towards significance (OR 4.6, [95% CI: 0.98-21.22], p=0.053).

**Table 3.**
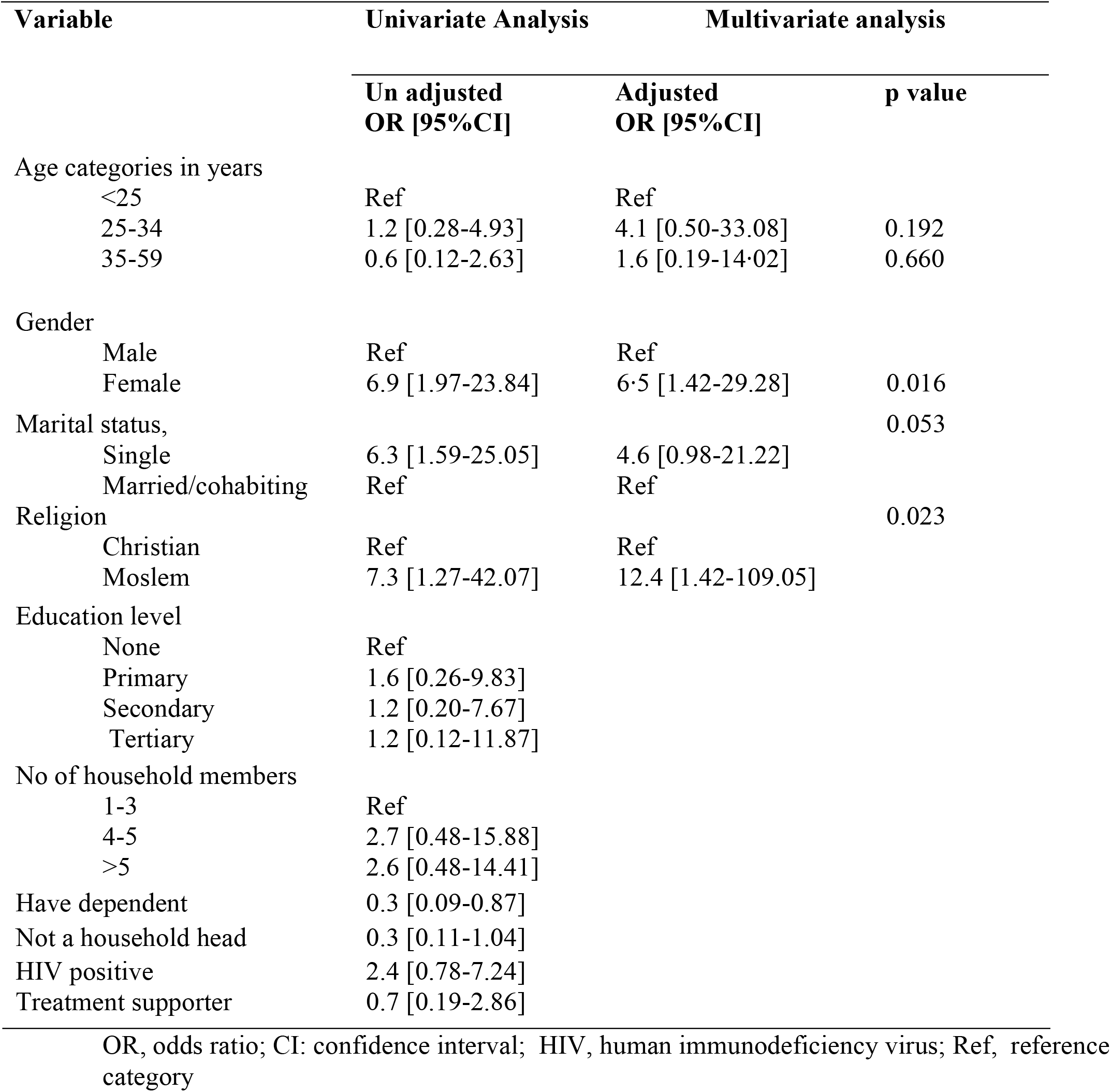
Results of univariate and multivariate analysis for factors associated with non-disclosure of adult TB status to household members

### TB patients’ positive post-disclosure experiences

Of the 57 patients with either complete or partial disclosure, 56 (98.3%) reported positive experiences post-disclosure. These were in form of support from household members in areas of: medication intake; encouragement; in feeding; and financial support to enable attendance of clinic days. The areas with the least reported support offered by household members were; infection control (2.4%) and counselling about TB as a curable disease (4.9%). Fig 1

**Fig 1.**
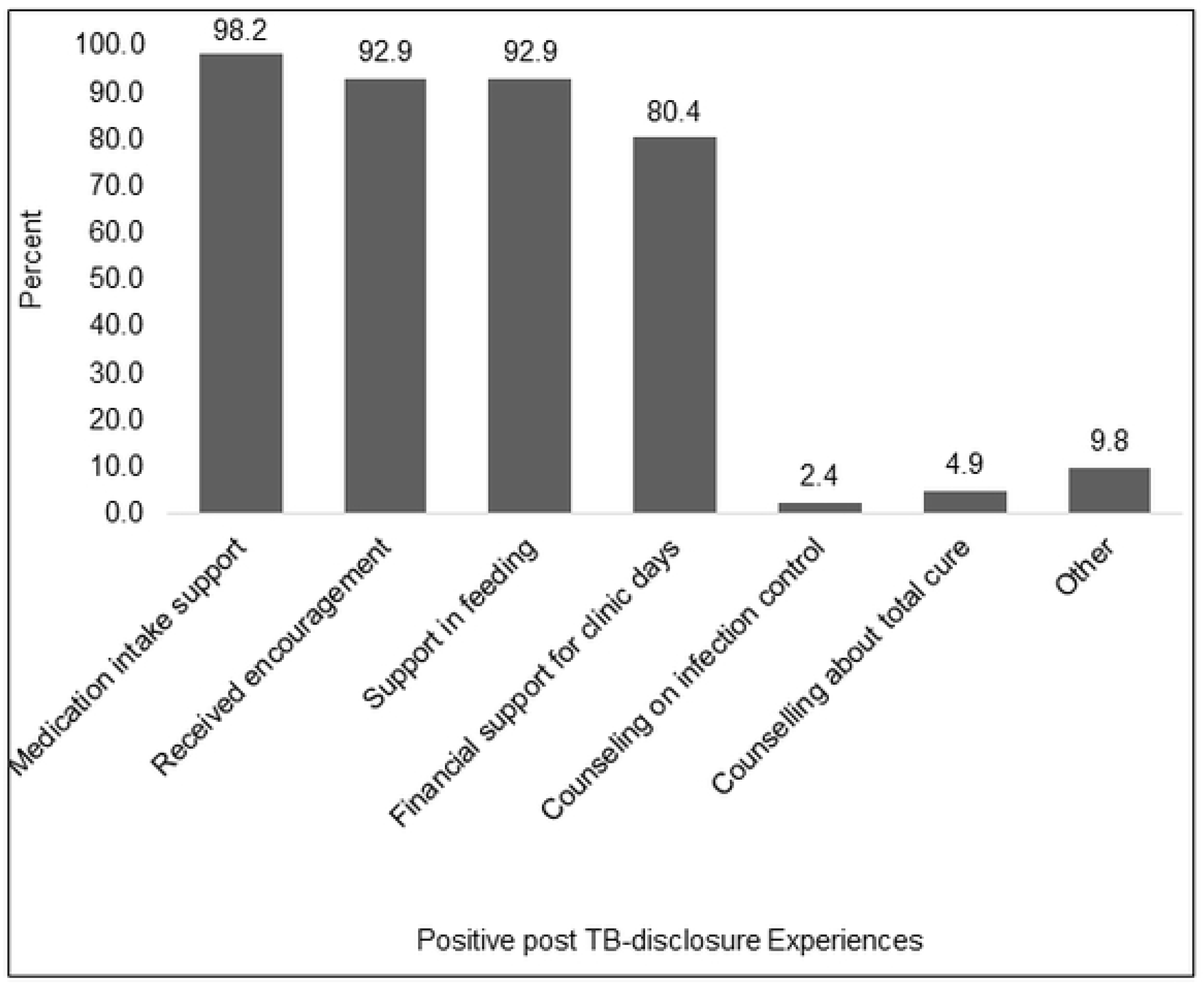
Positive post-disclosure experiences in adult TB patients, N=57

Majority of patients felt that these positive experiences post disclosure, had impacted greatly on their: treatment intake, attendance of clinic visits, self-esteem, hope of recovery, household harmony and work. Fig 2

**Fig 2.**
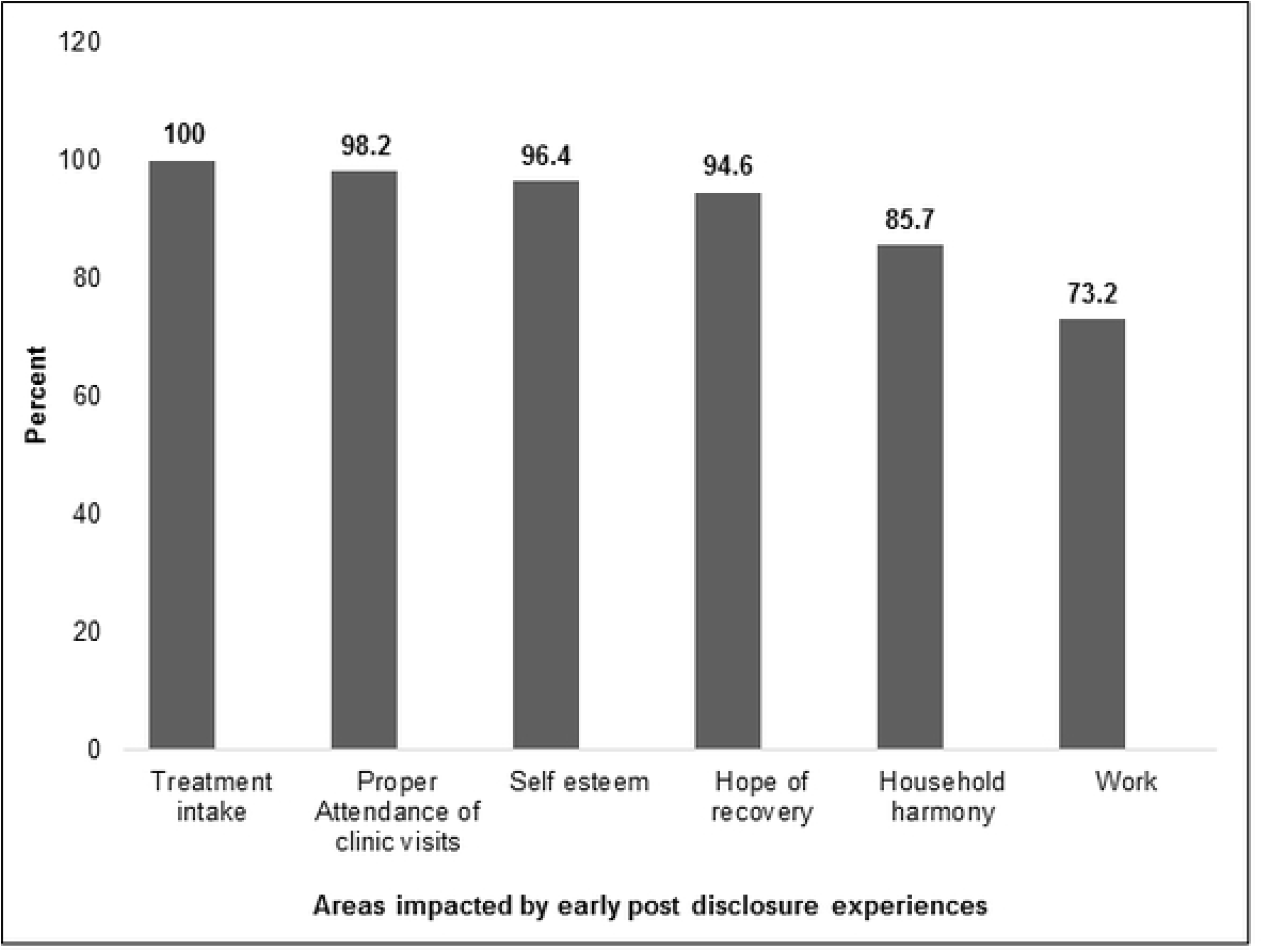
Areas of TB patients that are impacted by early positive post-disclosure experiences, N=56

### TB patients’ negative post-disclosure experiences

Of the 57 patients with either complete or partial disclosure, 14 (24.6%) had negative experiences following disclosure. The most common experience among them was thoughts of committing suicide (35.7%, n=5), marital separation (23%, n=3), blame and negative criticism (14%, n=2) and isolation (7%, n=1). Fig 3

**Fig 3.**
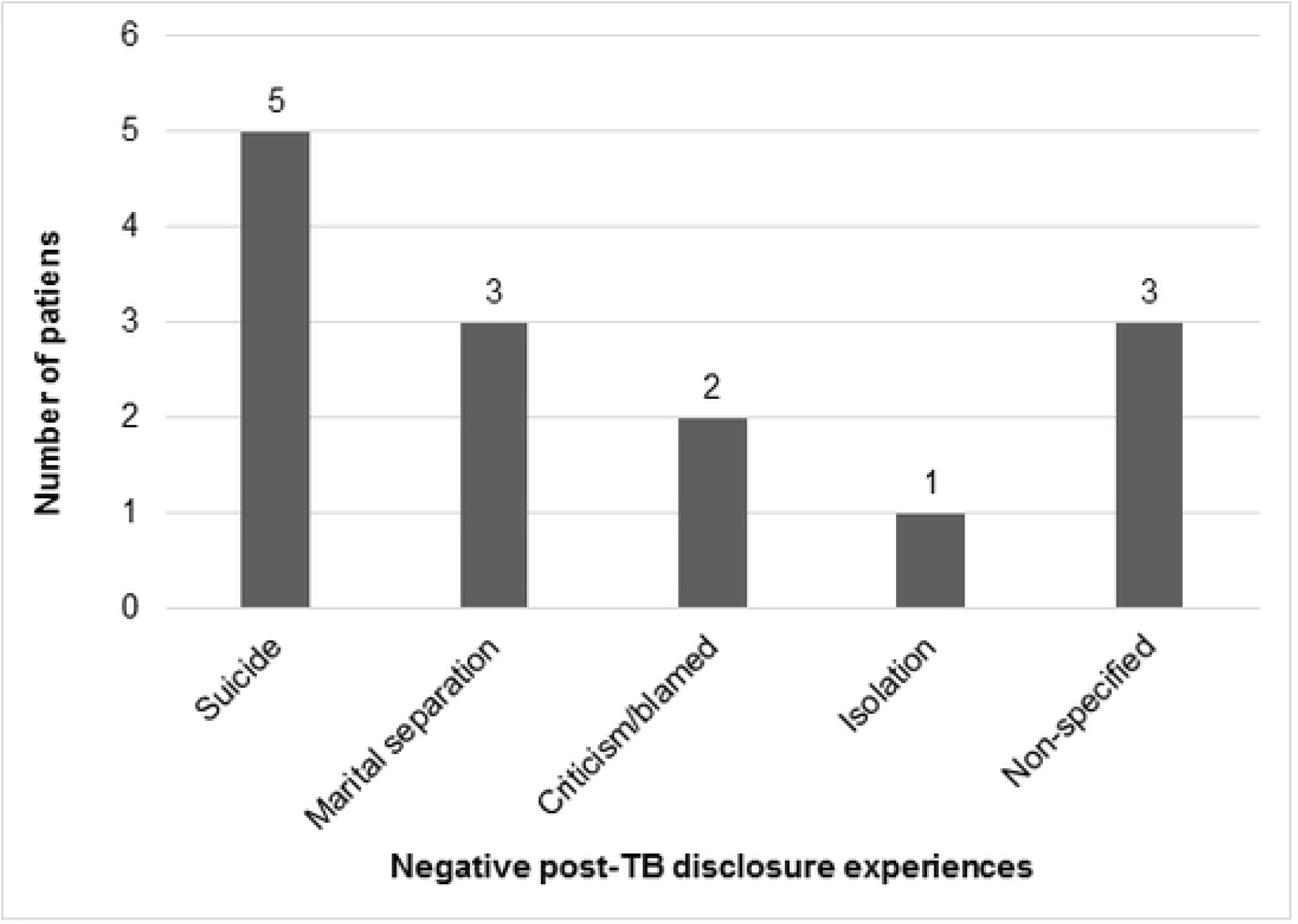
Negative post-disclosure experiences in adult TB patients, N=14

### Perceptions towards enhancing treatment supporter role in ensuring complete patient disclosure

All the 62 study participants whose perception on involvement of a treatment supporter to assist in patients’ TB disclosure to household members through training and provision of suitable TB information aides (like charts) by healthcare workers was explored, 38 (61.3%) expressed their agreement.

## DISCUSSION

Our study is the first to our knowledge to explore the dynamics of TB disclosure at a household level in Uganda, as a key strategy in promoting engagement of household members in TB control and efforts to achieve the targets of the End TB strategy [15]. We report a high rate of non-disclosure to household members by adult TB patients (30.6%). The need to attract care was the main motivational factor for patients to disclose to household members, besides other factors like: the desire to avoid transmitting the disease to them and attracting support in picking or reminding them to take their medicines. TB disclosure to household members yielded positive benefits almost to all patients, especially with regard to treatment adherence and psychosocial support, but with little gains on counselling pertaining to infection control and hope of complete cure from TB. Negative experiences occurred to one-in-four TB patients following disclosure to household members. Being female or a Muslim was significantly associated with non-disclosure of TB diagnosis to household members. A potential intervention based on training and provision of suitable TB information aides to treatment supporters by healthcare workers, so as to enhance their confidence in supporting the TB patient in the process of disclosure to household members so as, was acceptable by two-thirds of TB patients.

The observed high TB non-disclosure rate among adult patients to their household members, is of great public health concern, and can result in uncontrolled TB transmission at household level. Some of the patient factors and their household characteristics in our study seem to be in alignment with this concern, and indicative of a conducive environment for continued TB exposure and potential transmission. The point in case is the fact that; 1) majority (80%) of TB patients were staying with at least 4 household members, who would therefore qualify as TB contacts and at-risk of TB infection; 2) most of the patients (95%) were newly diagnosed with TB, and though not explored in this study, but could perhaps have had insufficient experience and knowledge on TB infection control measures so as to be able to minimize transmission to their household members; and 3) as part of the study eligibility criteria, all patients had been atleast 2 weeks on treatment, and we noted a median time of 13.3 weeks since treatment initiation among patients reporting non-disclosure to household members. This suggests a prolonged period of potential uncontrolled TB transmission to household members. Such non-disclosure deprives the patient with TB of the care and support they would have received from their household members. In addition, it deprives their household members especially the children or HIV infected contacts who are more likely to develop disease and to present with severe forms of TB once infected, of their right and early opportunity to self-protection and access to TB services. Such TB servicesinclude; contact screening with initiation on preventive treatment, prompt TB diagnosis, and early initiation on appropriate TB treatment. This is an important reason why the “ethical and human right approach” has been proposed to guide the pursuit of END-TB targets [18].

The finding in our study that disclosure was motivated by the patient’s desire to attract care from household members further emphasize the culturally perceived role of family in Africa in caring for their patients. Such a perception though good, positions household members at risk of contracting TB as they care for their patients if appropriate TB infection control measures are not promoted and enhanced at this level. On the other-hand, it could be expected that anything that could deny a patient this care and support from household/family members could potentially impact on their psychosocial status, treatment adherence, and outcomes of treatment. Therefore, our findings further support the need for social and psychological support interventions within TB patient care [7].

Our study showed that being a female adult TB patient was associated with 6.5 times higher odds of partial or total non-disclosure as compared to males. In as much as the reasons for this gender disparity were not explored in this study, one potential explanation would be on the underlying fear of TB-related stigma and discrimination that could result from disclosure of TB status, just as it has been reported in studies conducted in Zambia and Ghana, with great effect on women than men [13, 19]. Other fears for disclosure among female TB patients could be on the risk of breaking marital harmony [14]. Importantly, this disparity raises important concerns with a need to explore gender issues within the efforts tailored towards TB control or elimination. Given that the study participants were from a setting in which women rely on their husbands for financial support to access healthcare, non-disclosure could deny women of this financial support for drug collection and could result in poor adherence as already reported in the context of patients on ART in Africa [20]. Also, given the culturally perceived role of women, that is, to care for people within their households, positions them as important agents of TB transmission especially to those dependent on their care like under-five children, their spouses and others in vulnerable states e.g. the HIV positive, pregnant women and elderly. More to this, given that the women in rural Africa, bare the greatest burden pertaining to household food production, children feeding, food preparation and child healthcare [21], issues related to their health could impact greatly on their children’s health, especially whether they could be enrolled on preventive treatment or evaluated for TB in the timely manner.

Belonging to the Muslim faith had 12.4 times higher odds of non-disclosure of TB status to household members as compared to patients belonging to the Catholic faith (p=0.023). This is a new finding especially in the context of TB disclosure. However, shame-related HIV stigma has been reported to be strongly associated with religious beliefs such as the belief that HIV is a punishment from God or that people living with HIV/AIDS have not followed the Word of God [22]. It is however not clear whether such shame-related stigma, motivated by religious beliefs do exist in the context of TB, and if it could explain the disparity in religions with respect to disclosure to household members.

As expected, our study reports positive rewards for majority of patients (>98%) following TB disclosure to their household members. The commonest reported positive rewards include; household members supporting their medication intake, offering encouragement, supporting their feeding, and financial support to enable attendance of clinic days. As expected, such positive rewards of disclosure, were considered by patients to have impacted on their treatment adherence and psychosocial aspects. This finding affirms the benefit of a successful engagement of family/household members in TB patient care through patients’ TB disclosure, a known foundational strategy in ensuring good treatment outcomes and infection control [11].

Surprisingly, negative experiences following TB disclosure to household members were also noted to manifest concurrently with the positive experiences discussed already above. These occurred in one-in-three of TB patients that had disclosed their TB diagnosis to all or some of their household members. Such experiences especially; thoughts of committing suicide (35.7%) and marital separation (23%), but also blame and negative criticism, and isolation, seem to indicate the existence of severe effects of potential underlying TB-related stigma among household members and communities at large. These findings are in agreement with those reported in previous studies on patients’ TB disclosure to household members, that reported isolation, divorce, among other outcomes [14]. Such negative post-disclosure experiences could be explained by the low awareness about TB disease that could be existing among patients and household members, something that could complicate TB control [23]. Such deficit in knowledge on TB by household members of TB patients, could be represented indirectly in this study by the fact that counseling on infection control and hope for total cure from TB, where the least performed aspects by household members following patients’ TB disclosure. Such deficit in TB knowledge, could offer alternative explanation on why negative experiences erupted concurrently with positive experiences of patients post disclosure. These findings, not only highlight the need for robust interventions aimed at enhancing social and psychological support to TB patients [7] through TB awareness, but also interventions that could directly promote TB disclosure to household members while minimizing on such negative outcomes.

Enhancing the involvement of treatment supporters in facilitating successful stigma-free disclosure of TB by patients to household members through training and provision of suitable TB information aides by healthcare workers, is one potential intervention that was perceived acceptable to two-thirds of TB patients in the study. Since the treatment supporter system is already in existence to support home-based DOTS, and that all TB patients are required to have one where necessary prior to treatment initiation, such disclosure enhancement interventions centered on treatment supporters could easily be adopted. However, the feasibility and efficacy of such a models would need to be evaluated in future randomized trials, and its success would depend largely on the healthcare workers’ level of knowledge of TB, an argument supported by the already known positive correlation in knowledge on infection control practices between healthcare workers and household members [24]. Efficacy would also depend on the availability of acceptable TB information aides for use by both patient and treatment supporter, so as to facilitate the disclosure process to household members.

The study has several limitations: i) The small sample size may have reduced the power to detect significance of some predictor variables of non-disclosure. ii) The study did not quantify the level of stigma among patients, in as much as it is known to influence disclosure. iii) Patients’ disclosure or not of TB to the household members was not assessed in regards to some clinical or treatment outcomes such as completion, cure, adherence or household contact screening, something that should be evaluated in future studies utilizing stronger analytical study designs. iv) The fact that the study never utilized qualitative assessment methods could not allow complete understanding of the major hindrances of disclosure especially among the female and moslem patients.

On the other-hand, our study unfolds the less addressed problem of TB patient non-disclosure to household members that could potentially impend the success of TB control strategies and attainment of the End TB targets of reducing TB incidence and TB-related death by 95% by 2035 [15]. Some of such TB control strategies hinged on successful disclosure to household members are: 1) TB Contact screening, which is impossible unless an index case voluntarily reports to have household contacts and willing to disclose to household members, 2) Initiation of TB contacts on preventive treatment, especially children, HIV infected adults, and other vulnerable household members, 3) Early TB diagnosis among symptomatic household members, which may be limited by delayed TB suspicion, hence delayed health seeking process by household members of TB patients, 4) Implementation of appropriate infection control measures at household level, 5) Home-based DOTS, and 6) Involvement of household members to offer social and psychological support to their patients, something that would impact on their treatment adherence and outcomes.

### Recommendations

Based on the study findings, we recommend that: 1) future larger studies should be conducted to conclusively establish the predictors of TB non-disclosure to household members, post-disclosure experiences, and their impact on treatment outcomes within both rural and urban settings; 2) Development of TB information aides to facilitate in process of patient disclosure to household members should be prioritized; 3) Interventions for enhancing disclosure at household level that are gender and religion-sensitive should be developed and evaluated in well-designed community-based randomized trials so as to ascertain their efficacy and impact on ensuring successful stigma-free TB disclosure, early access to TB diagnostic services, preventive therapy uptake, treatment adherence and extent of family support to the TB patients; 4) Psychosocial counselling should be integrated within the healthcare package for TB patients among TB burden countries; 5) Qualitative studies are needed to explore the reasons for non-disclosure among female and Muslim TB patients.

### Conclusion

The rate of non-or partial TB disclosure to household members is worryingly high, with important disparities across gender and religion. This could risk missing an opportunity to control TB transmission at household level, access to available TB diagnostic, treatment and preventive services by household members, and attracting family support towards their patients’ care. Negative post-disclosure experiences reflecting severe forms of TB-related stigma, co-exist amidst the enormous patient-felt benefits of disclosure. There is need to enhance the knowledge of household members and patients on TB infection control measures so as to minimize TB transmission at household level. Gender and religion sensitive interventions to enhance TB disclosure at household level while minimizing effects of stigma are urgently needed so as to increase engagement of household members in TB control efforts and patient care.

## DECLARATIONS

### List of abbreviations

(TB): Tuberculosis;
(DOTS): Directly Observed Treatment, Short-course;
(HIV): Human Immunodeficiency Virus;
(AIDS): Acquired Immunodeficiency Syndrome;
(OR): Odds Ratio;
(CI): Confidence Interval.

### Consent for Publication

Not applicable

### Availability of data and materials

The dataset from which the information presented in this manuscript originates has been submitted as additional supporting file.

### Authors’ Contributions

DA MN conceived the study, and participated in its design. MN DA and FB participated in data collection, study monitoring and coordination of the study. DA MN FB performed the statistical analysis. DA MN FB drafted the manuscript. All authors read and approved the final manuscript.

## Acknowledgments

The authors wish to appreciate Dr Maryline Bonnet of IRD UMI233/ INSERM U1175, France, for the critical review of the manuscript. Special appreciation to the study participants, the valuable contributions of the health care workers at TB clinic-Mbarara Regional Referral Hospital. We are grateful to Arimpa Amerias and Kyarimpa Rose for the assistance during data collection.

